# Live imaging and functional characterization of the avian hypoblast redefine the mechanisms of primitive streak induction

**DOI:** 10.1101/2025.05.15.654239

**Authors:** Aurélien Villedieu, Olinda Alegria-Prévot, Carole Phan, Yu Ieda, Francis Corson, Jérôme Gros

## Abstract

In birds and mammals, the formation of the primitive streak, the hallmark of the primary axis and site of gastrulation, is thought to occur when an anterior displacement of the hypoblast (visceral endoderm in mice) lifts its inhibition on the posterior epiblast, enabling the activation of NODAL signaling. Although the anterior movement of the murine visceral endoderm is well documented, the dynamics of the avian hypoblast remain poorly understood. Here, using live imaging and quantitative image analysis, we find that the hypoblast is mechanically coupled to the epiblast and does not migrate away from its posterior end. Instead, the hypoblast moves and deforms passively, in response to the forces transmitted from the epiblast that shape the primitive streak, after its induction. Furthermore, we show that the posterior hypoblast does not exert an inhibitory effect on the epiblast but instead expresses *NODAL*, which activates primitive streak formation. NODAL concomitantly regulates gene expression in the hypoblast, patterning it along the anteroposterior axis. Our results thus redefine how the primary axis is established in avians, demonstrating that the displacement of the hypoblast and its concomitant anteroposterior patterning are consequences *—* rather than drivers *—* of primitive streak induction, downstream of NODAL signaling.

**Essential summary:** *NODAL* asymmetric expression in the hypoblast activates primitive streak formation in the avian embryo.

## Introduction

In amniotes, the basal side of the epiblast is covered by a layer of extra-embryonic endodermal cells called the hypoblast (Figure 1A, top). The hypoblast plays a critical role in inducing the primitive streak, which defines the embryonic primary axis. In mice and rabbits, the hypoblast (called visceral endoderm [VE] in mice) secretes inhibitors of NODAL and WNT signaling pathways (e.g. CER1, LEFTY1 and DKK1) that inhibit primitive streak formation in the epiblast^1,2^. In mice, this inhibition is initially located in the distal part of the VE and is then relocated to the anterior side of the embryo owing to an active migration of the VE^3^. This anterior movement of the VE, whose direction has been proposed to be predetermined by early molecular asymmetries in the VE^4,5,6^, ensures the definitive positioning of the primitive streak at the posterior end of the epiblast and prevents the formation of additional primitive streak^1,7^.

**Figure 1:**
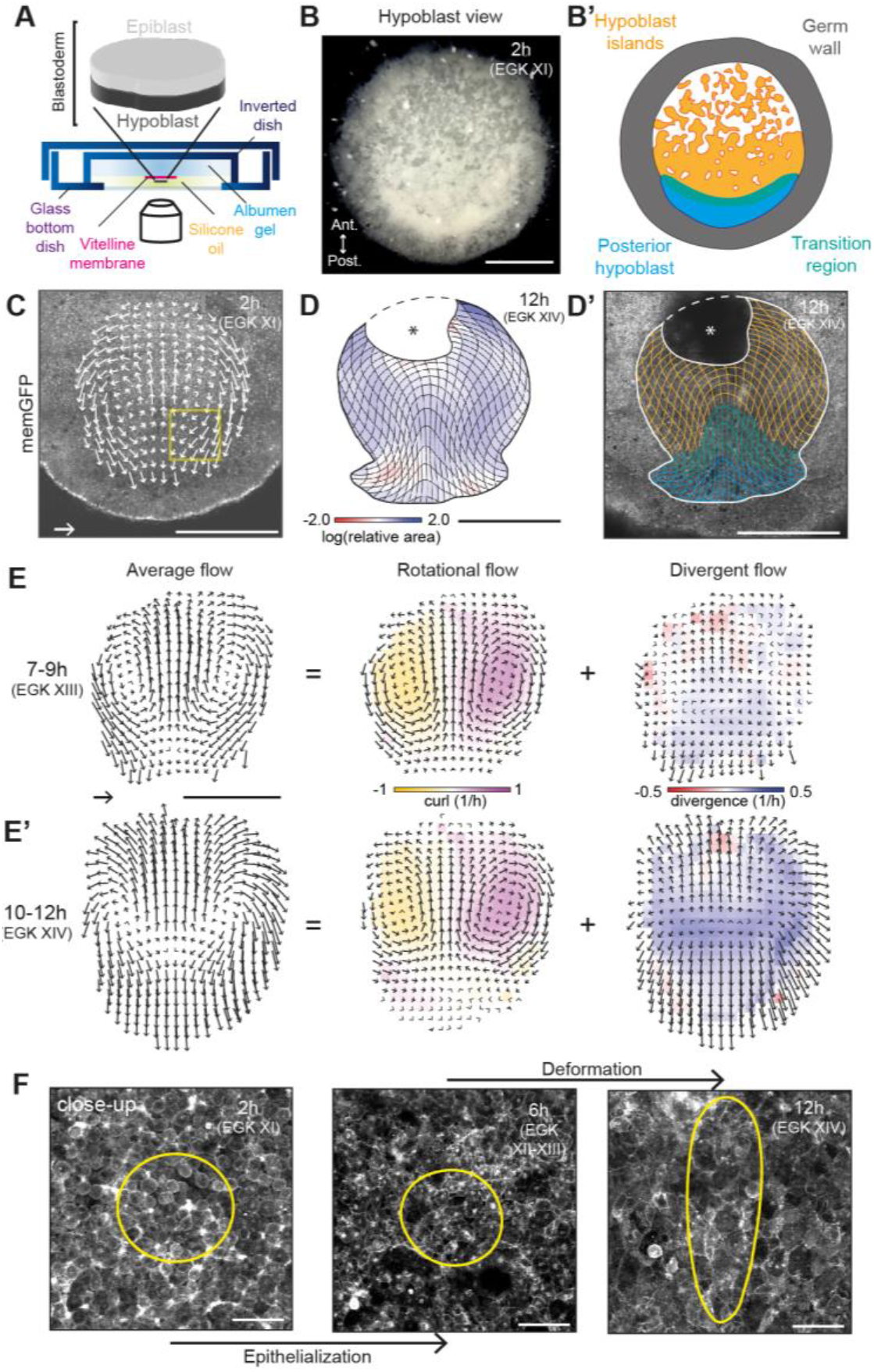
Counter-rotating flows shape the hypoblast. **(A)** Schematic of the *ex ovo* culture system used for high-resolution live imaging of the hypoblast. **(B)** Quail blastoderm viewed from the hypoblast side at 2h post-laying, and **(B’)** associated schematic showing the different hypoblast populations. The transition region joining the hypoblast islands (in orange) and the posterior hypoblast (in blue) is represented in green. **(C)** First image of a time-lapse imaging of the hypoblast in a memGFP transgenic embryo (2h post-laying). Arrows show the averaged hypoblast flows between 2h and 12h. The yellow square marks the location shown in closeup in Figure 1F. **(D)** Cumulated deformation of an initially squared grid (computed from PIV-calculated velocity fields) between 2h-12h, color-coded for area changes and overlaid on the last timeframe of the time-lapse experiment (12h) **(D’)** The grid is colored according to populations pictured in Figure 1B’. Asterisk: region masked by debris, in which PIV calculation is not reliable. **(E-E’)** Averaged velocity field within the hypoblast between 7-9h and between 10-12h (average of 9 embryos) and corresponding decomposition into rotational and divergent components of the flow. **(F)** High magnification time series of the yellow region shown in (C) at 2h, 6h and 12h. Yellow contours: region tracked by PIV. Scale bars: 1 mm (C-E’), 100 µm (D). Scale vectors: 100 µm/h.

In avian embryos, the site of primitive streak formation has also been proposed to result from interactions with the hypoblast, although its role has long been debated^8,9,10,11,12,13,14^. The hypoblast had initially been proposed to act as an inducer of primitive streak formation^8,11,15^, but this view was later challenged. Based on transmitted light time-lapse videomicroscopy and graft experiments, it has been proposed that the hypoblast is displaced anteriorly by the emergence of another extra-embryonic endoderm population, the junctional endoblast (also named secondary hypoblast), at the posterior end of the blastoderm^16,17,18,19^ (stages EGK XIII-XIV). The existence of this endoblast population and its role in displacing the hypoblast found support through the observation that the expression of genes in the hypoblast (e.g. *FOXA2, HHEX, CRESCENT*, etc.) including NODAL and WNT inhibitors (e.g. *CER1* and *DKK1*) becomes progressively relocated to the anterior end of the hypoblast, as the primitive streak appears in the posterior epiblast^13,20,21,22,23^. From these observations, it was concluded that the emergence of the endoblast, by displacing the primary hypoblast anteriorly, relieves the posterior epiblast from inhibitors secreted by the hypoblast, entailing the formation of the primitive streak^13,16^. An inhibitory role of the hypoblast in primitive streak formation is supported by experiments involving its surgical ablation and reported to induce supernumerary axes — albeit at low frequencies^13^. Thus, as in mice, an anterior movement of the hypoblast relative to the epiblast would be a critical step in instructing primitive streak formation and positioning^1,24,25^. However, this model lacks critical experimental support in avians. While original studies have described an overall anterior motion of the hypoblast^17,19,20,26,27^, these are based on poorly resolved transmitted light imaging or experiments in which group of cells are sparsely labelled and followed using vital dye or carbon particles. Furthermore, despite the central role of a hypoblast motion relative to the epiblast in the proposed “relief-of-inhibition” model for primitive streak induction, such relative motion has not been directly observed. Indeed, counter-rotating tissue flows accompanying primitive streak formation occur simultaneously in the epiblast^14,28,29^, and how they compare with the tissue flows of the hypoblast is not known. Finally, it is unclear whether the anterior relocation of gene expression within the hypoblast results from a physical displacement or from genetic regulation within the hypoblast itself.

In this study, we characterize the dynamics of the avian hypoblast and show that it is passively displaced by the forces driving primitive streak formation in the epiblast. Thus, the posterior hypoblast remains in constant contact with the presumptive primitive streak territory of the epiblast, as gastrulation movements take place. These findings which are inconsistent with a relief of inhibition prompted us to re-evaluate the inhibitory role of the hypoblast in controlling primitive streak formation. At odds with earlier proposal, we find that *NODAL* expression initiates in the posterior hypoblast and then propagates to the epiblast, where it induces the formation of the primitive streak while patterning the hypoblast along the anteroposterior axis. Collectively, our findings, which clarify the dynamics and role of the hypoblast, redefine the cellular and molecular mechanisms underlying primary axis induction in avians.

## Results

### Counter-rotating flows shape the hypoblast

To capture the dynamics of the hypoblast, we first developed a culture system for high-resolution imaging of transgenic quail embryos expressing fluorescent reporters (Figure 1A). In this set-up, the Early Chick (EC) culture system^30^ is inverted onto a glass-bottom dish, and silicone oil is placed between the hypoblast and coverslip (see Methods). Using this system, hypoblast development could be imaged from 2h to 12h after egg-laying (corresponding to EGK XI-XIV^31^) in 9 embryos expressing a membrane-bound green fluorescent protein^29^ (memGFP; Figure 1B-F, Movie 1). These movies provide a cell-resolved description of the developmental stages defined macroscopically by Eyal-Giladi and Kochav^31^. In line with their observations, we observe that at the laying stage (stage EGK XI), the avian hypoblast consists of isolated non-epithelialized islands in its center, joined by a denser, opaque posterior hypoblast crescent known as Koller’s or Rauber’s sickle^15^, surrounded by a thicker peripheral ring of another extra-embryonic tissue called the germ wall^16,31^ (Figure 1B-B’, Supplementary Figure 1A). This organization is confirmed by immunofluorescence against FOXA2 (also called HNF-3β), which is expressed throughout the hypoblast at this stage^22^ (Supplementary Figure 1B). At around 5h, the hypoblast, which initially comprised rounded cells, start to fuse to form a cohesive epithelial tissue (Stage EGK XIII^31^; Figure 1F, Movie 1). The cohesiveness of the hypoblast from 7h can also be observed when an attempt is made to separate the epiblast from the hypoblast by microsurgery: while it is difficult to remove hypoblast islands at 2-5h, it becomes easy to dissect the hypoblast in one piece after 7h (Supplementary Figure 1C).

The motion of the hypoblast can be quantified over time using Particle Image Velocimetry (PIV; see Methods; Figure 1C-F, Movie 1). Between 2-6h, the initially rounded cells of the hypoblast epithelialize *in situ* without major tissue movement, as revealed by following PIV-tracked regions (Figure 1F and Movie 1). From 5h onwards, the hypoblast shows large-scale tissue flows. Surprisingly, we find that the hypoblast does not display a strict anterior migration but exhibits counter-rotating flows (Figure 1C, 1E-E’), very similar to the ones that accompany primitive streak formation in the epiblast^29,32^. As a result, the shape of the primitive streak can be recognized in the posterior hypoblast, which converges along the medio-lateral axis and extends along the anteroposterior axis (Figure 1D-F). However, several differences remain compared to the morphogenesis of the primitive streak in the posterior epiblast. First, the posterior hypoblast expands as it converges and elongates (Figure 1D-E’), in contrast to the extensive tissue contraction that characterizes the crescent-shaped presumptive primitive streak territory in the epiblast^29,32^. Second, cells of the posterior hypoblast, facing the primitive streak, elongate along the anteroposterior axis (Figure 1F, Movie 1), in contrast to the mediolateral elongation of cells in the epiblast evidencing an actively driven convergent extension of the primitive streak. Decomposing the flows into rotational (incompressible) and divergent (area changes) components^29^ reveals the persistence of counter-rotating flows throughout the 10 hours movie and identifies an area expansion flow that is maximal from 10 to 12h (Figure 1E-E’). To ensure that the PIV calculation of the hypoblast flows is not mistaken by integrating fluorescence from epiblast cells, the hypoblast of wild type (WT) embryo was surgically replaced by a hypoblast expressing a H2b-GFP reporter gene^33^ at 7h (Supplementary Figure 2A-B). The resulting chimeras, in which only the hypoblast cells express H2b-GFP, confirmed the presence of the above-mentioned counter-rotating flows and tissue deformation (Supplementary Figure 2C-D, Movie 2).

### The epiblast entrains and deforms the hypoblast during primitive streak formation

To characterize the motion of the hypoblast relative to the epiblast, we photoconverted small regions of transgenic embryos expressing the green-to-red photoconvertible mEOS2 protein^34^ at 2h. Because of the low Z-resolution of confocal excitation, both epiblast and hypoblast cells were photoconverted in these regions, which were then imaged by orientating the blastoderm on its hypoblast or epiblast side (see Methods). Despite extensive tissue movement, 6 hours later, photoconverted cells in the hypoblast and epiblast remained in close association, although cells from hypoblast regions were more dispersed than their epiblast counterparts (Figure 2A, Movie 3). Thus, the hypoblast and epiblast display qualitatively similar counter-rotating flows, and the two layers show little relative displacement. To be more quantitative, we grafted early hypoblast islands expressing tdTomato:Myosin^29^ onto a memGFP-expressing host at 2h, prior to hypoblast epithelialization, so that grafted cells could be incorporated during the epithelialization of the hypoblast, and we simultaneously live imaged the epiblast and hypoblast dynamics thanks to the differential expression of tdTomato:Myosin and memGFP (Figure 2B, Movie 4). Quantification of the epiblast and hypoblast flows show that they are highly correlated, confirming results obtained with mEOS2 photoconversion (Figure 2C, Supplementary Figure 3A-C). Subtracting the velocity fields of the hypoblast from those of the epiblast (Figure 2C’, Supplementary Figure 3D) reveals that the relative displacement between these tissue layers is small compared to the overall displacement of the epiblast and hypoblast (compare Figures 2C and 2C’) and corresponds mostly to an area expansion flow (Figure 2D-D’). Altogether, these experiments suggested that the rotational component of the flow observed in the hypoblast might result from mechanical coupling with the epiblast, in which active stresses at the embryo margin produce the large-scale rotational tissue flows associated with primitive streak formation^29^. To test this hypothesis, a porous filter was inserted between the hypoblast and the epiblast at 7h (Figure 2E). If diffusible molecules can be exchanged through this filter, mechanical contact between the two layers is however prevented^35^. In this condition, tissue flows occurred normally in the epiblast, resulting in primitive streak formation as in control embryos in which the hypoblast was dissected and put back in place without inserting a filter (Figure 2F-F’); the hypoblast however did not exhibit counter-rotating flows, unlike control embryos, but only exhibited a small area expansion (n=7/7 with filter, n=4/4 controls, Figure 2G-G’, Movie 5). These results demonstrate that mechanical coupling between the epiblast and hypoblast is required for the rotational tissue flows to occur in the hypoblast, whereas they proceed autonomously in the epiblast. They further show that these flows arise from the passive transmission of active forces generated in the epiblast while only an area expansion is intrinsic to the hypoblast. Consistent with this, we find that the hypoblast shows no detectable levels of junctional phosphorylated Myosin II (which is a read out of active force generation), in contrast to the epiblast, where supracellular cables of phosphorylated Myosin II could be readily identified at the margin of the same embryo, as previously published^29^ (Supplementary Figure 2E). Altogether, these results show that the tissue flows observed in the hypoblast can be understood as the sum of a passive rotational flow entrained by the active forces generated in the epiblast owing to mechanical coupling and an active but modest expansion flow intrinsic to the hypoblast layer (Figure 2H).

**Figure 2:**
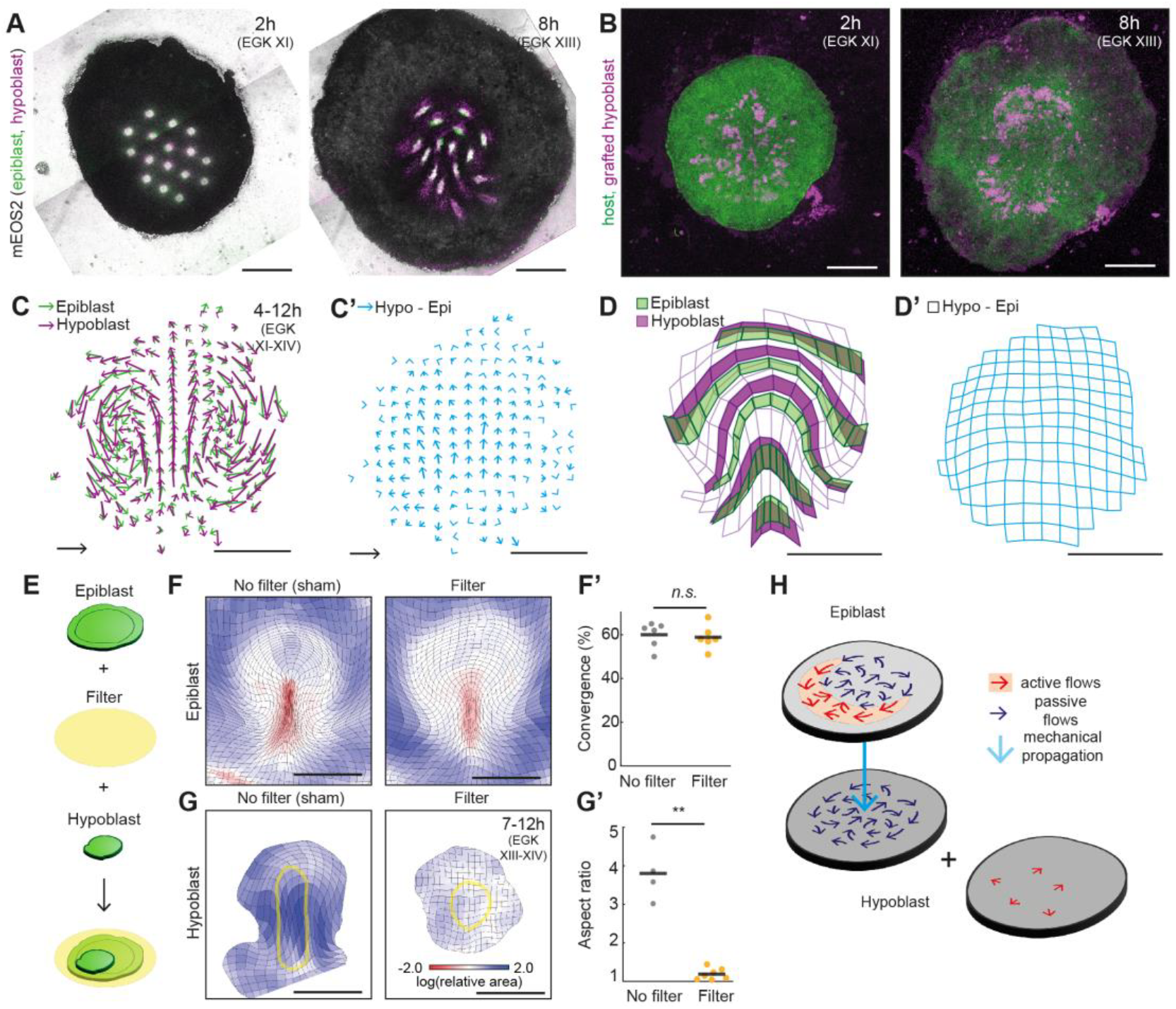
The epiblast entrains and deforms the hypoblast during primitive streak formation. **(A)** mEOS2 transgenic embryo immediately after (left) and 6h after (right) the photoconversion of 16 square regions. Photoconverted cells of the hypoblast and of the epiblast are shown in magenta and green, respectively (in n=7/7 embryos, photoconverted hypoblast and epiblast regions show little relative displacement 6h after photoconversion). **(B)** memGFP transgenic embryo (green) grafted with hypoblast cells expressing tdTomato:Myosin (magenta) at 2h and 12h. **(C)** Averaged epiblast and hypoblast flows between 4h-12h (from n=10 grafted embryos). **(C’)** Averaged difference between hypoblast and epiblast flows shown in (C). **(D)** Cumulative deformations of the hypoblast (grid in magenta) between 4h-12h. 5 initially horizontal stripes of the grid are shown in plain magenta and corresponding stripes in the epiblast are shown in plain green, to visualize the relative deformation between the two tissue layers. **(D’)** Grid representing the averaged deformation of the hypoblast after the deformation of the epiblast has been subtracted. **(E)** Sketch illustrating the intercalation of a porous filter between hypoblast and epiblast at 7h post-laying. **(F, G)** Grid showing the cumulative deformation between 7h-12h of the epiblast (F) and the hypoblast (G) with (bottom) or without (top) filter intercalation. A circular region (yellow circle) in the center of the hypoblast at 7h is tracked by PIV in (G). Color code represents area changes. **(F’, G’)** Quantification of the medio-lateral convergence of the primitive streak in the epiblast (F’) and aspect ratio of PIV-tracked regions in the hypoblast (G’) at 12h with or without intercalation of a porous filter (F’, n=6 embryos with or without filter, *n*.*s*.: non-significant, Welch test p-value = 0.73; G’, n=4 and 7, **: p-value<0.001, Welch test p-value=0.005). **(H)** Diagram of the hypoblast motion as the sum of a passive reaction to the contractile forces generated in the epiblast and a small active deformation (red: active flows, blue: passive flows, light blue arrow: propagation by mechanical coupling). Scale bars: 1 mm. Scale vectors: 100 µm/h.

### *NODAL* expression initiates in the posterior hypoblast and propagates in correlation with the progressive patterning of the hypoblast

Our results show that there is little relative displacement between the hypoblast and the epiblast at the onset of gastrulation. Thus, the hypoblast remains in constant contact with the primitive streak during its formation, an observation which is inconsistent with an inhibitory function of the hypoblast on primitive streak induction. We thus decided to reanalyze the expression of *NODAL* and *GDF1*, two key inducers of the primitive streak^16,36^, using Hairpin Chain Reaction-based RNA fluorescent in situ hybridization (HCR-RNA-FISH). At odds with earlier reports^23,37,38^, we found that *NODAL* is first expressed in the posterior hypoblast (corresponding to the Koller’s sickle) at 2h (EGK XI), before the epiblast expresses *NODAL* or *GDF1*, in both quail and chicken embryos (Figure 3A-B, Supplementary Figure 4). To characterize the temporal evolution of *NODAL* expression, we labeled quail embryos between 2 hours (EGK XI) and 8 hours (EGK XIII), timing them precisely based on the progression of their gastrulation movements (see Methods). By spatially aligning and averaging fluorescent signals of 6-11 embryos per timing (see Methods), we generated archetypal maps of *NODAL* RNA levels over time (Figure 3B). These maps reveal that *NODAL* expression in the hypoblast increased over time and then appeared in the posterior epiblast by 4h, reaching expression levels comparable to those of the hypoblast by 6h. Thus, a wave of *NODAL* expression, initiating from the posterior hypoblast, propagates across the epiblast and hypoblast, forming an anteroposterior gradient in both tissue layers. *GDF1* expression, on the other hand, was detected later than *NODAL*, at around 5h (EGK XII) and specifically in the epiblast (Supplementary Figure 4). As observed for *NODAL, GDF1* expression increases over time in the epiblast and then appears in the hypoblast, but in a more anterior pattern than *NODAL*. These results suggest that *GDF1* is downstream of *NODAL* during primitive streak induction, although it remains possible that the HCR-RNA-FISH probes used here do not allow detection of very low expression levels.

**Figure 3:**
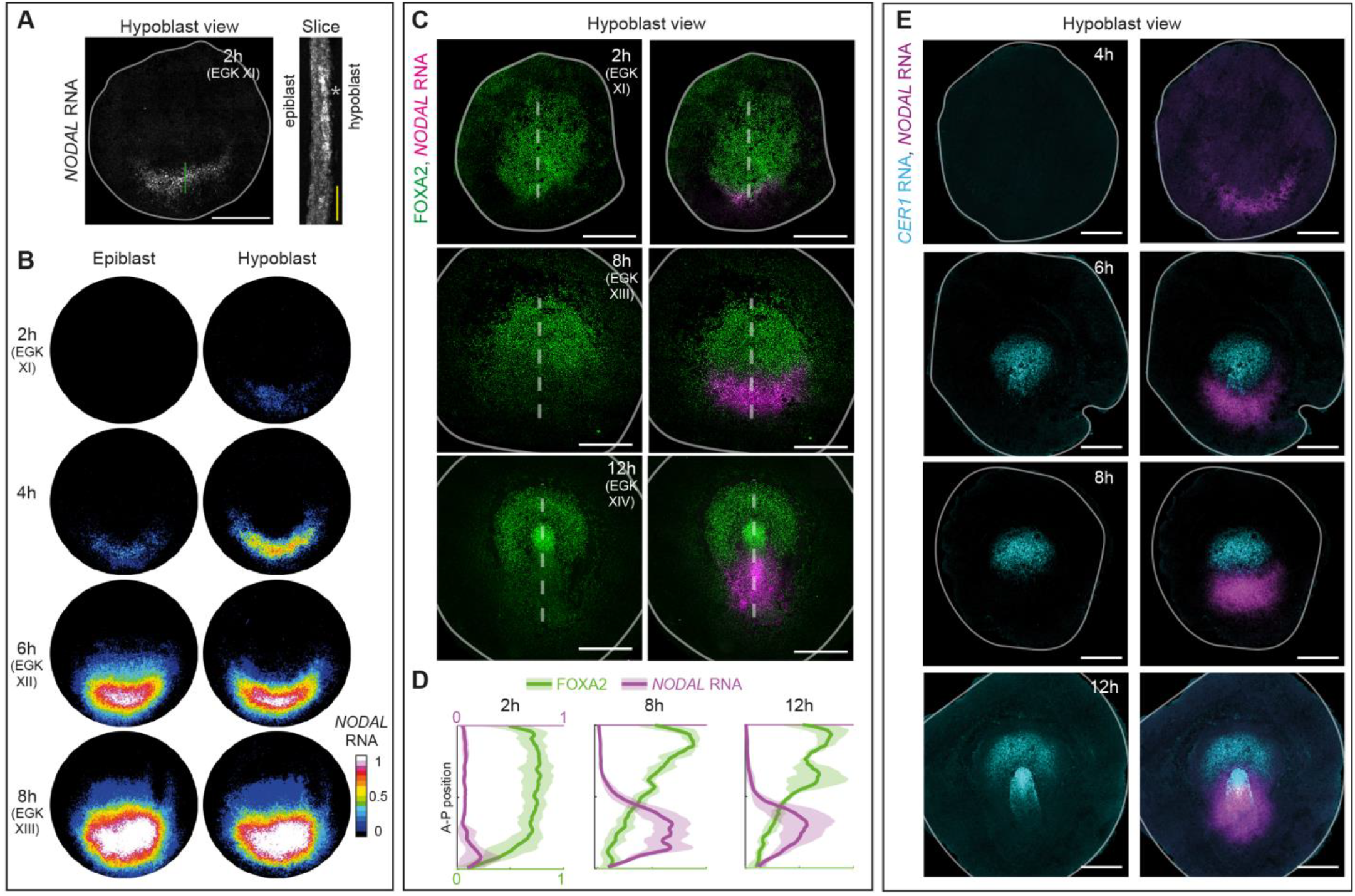
*NODAL* expression initiates in the posterior hypoblast and propagates in correlation with the progressive patterning of the hypoblast. **(A)** *NODAL* RNA levels at 2h post-laying. Left: hypoblast view, right: cryosection of the posterior zone (location of the section is marked by a yellow line). *NODAL* is expressed specifically in the hypoblast in n=3/3 sectioned embryos. **(B)** Archetypal map of *NODAL* RNA levels in hypoblast and epiblast at 2h (average of 7 embryos), 4h (average of 6 embryos), 6h (average of 11 embryos) and 8h (average of 7 embryos, see Methods). **(C)** Immunofluorescence for FOXA2 and FISH of *NODAL* RNA at 2h, 8h and 12h. **(D)** Average anteroposterior profiles of FOXA2 and *NODAL* RNA levels at 2h (average of 7 embryos +/-std), 8h (average of 7 embryos +/-std) and 12h (average of 4 embryos +/-std). **(E)** FISH of *CER1* and *NODAL* RNA at 4h, 6h, 8h and 12h. Scale bars: 100 µm (A right), 1 mm (A left, C).

We then investigated how this *NODAL* expression gradient relates to the antero-posterior patterning of the hypoblast. We first characterized the pattern of FOXA2 in relation to *NODAL* expression. As previously published^21,22^, we observed an initially homogeneous localization of FOXA2 in the hypoblast at 2h, that over the course of 12h progressively took the shape of a gradient, decaying from anterior to posterior (EGK XIV, Figure 3C left). Notably, this anteroposterior gradient of FOXA2 is anticorrelated with the gradient of *NODAL* expression (Figure 3C-D). As mentioned above, this anterior relocation of *FOXA2* expression (along with other genes) has been attributed to the displacement of the hypoblast by the invading endoblast^13,16,21^. However, backtracking the *NODAL*-high/FOXA2-low domain from 12h to 2h, using hypoblast tissue flows from the same embryo, maps this region to the transition between the posterior hypoblast and the central hypoblast islands, where FOXA2 is initially high and *NODAL* expression emerges (Supplementary Figure 5). We also analyzed the pattern of *CER1* expression in relation to the one of *NODAL*. Surprisingly, we observe very low, if any, *CER1* expression early on and found that *CER1* starts to be reliably detected in the anterior hypoblast from 6h, in a pattern which is, like FOXA2, opposite to *NODAL* expression (Figure 3E). Altogether these analyses argue that the anterior wave of *FOXA2* and *CER1* expression results from a genetic regulation in the hypoblast that is concomitant to the propagation of the wave of *NODAL* expression, rather than from the emergence of a new cell population (endoblast) displacing the hypoblast.

### NODAL signaling in the posterior hypoblast induces the primitive streak

NODAL activity is well known to induce ectopic primitive streak formation^13,39^, strongly suggesting that the posterior hypoblast is indeed an inducer rather than an inhibitor of primary axis formation. To test this, we surgically removed hypoblast cells (both the posterior hypoblast and hypoblast islands) of quail embryos at 2h (Supplementary Figure 6A). In 7/9 cases, this prevented primitive streak formation and *NODAL* expression, while the epiblast expanded extensively, presumably as the result of tension-induced stretching arising during epiboly^40^, which is unaffected by hypoblast surgical ablation (Figure 4A, Movie 7). In the remaining 2/9 cases, a reduced primitive streak formed, probably due to incomplete ablation of the posterior hypoblast (Supplementary Figure 6B). Similar results were obtained when the hypoblast was removed in chicken embryos incubated for 2h (Supplementary Figure 6C). Interestingly, an intentional incomplete removal of the hypoblast, leaving two pieces of posterior hypoblast on each side of the ablated zone (Supplementary Figure 6A) resulted in the formation of two primitive streaks facing the untouched hypoblast in 7/11 cases (Figure 4A, Movie 7), or a single primitive streak but shifted from its original site in the remaining 4/11 cases (Supplementary Figure 6B). These results not only show that, in the absence of the hypoblast, the primitive streak is not induced, but they also provide a possible explanation for the previously proposed inhibitory role of the hypoblast^13^. That proposal was based on supernumerary axes induction upon surgical ablation of the hypoblast *—* an outcome that could also be explained by incomplete ablation. Notably, the ablation of the hypoblast at later stages (7h) had no noticeable effect on primitive streak formation and *NODAL* expression, whether it was performed in quail or in chicken embryos (Supplementary Figure 6D), suggesting that the posterior hypoblast is required for primitive streak induction but dispensable for its maintenance. Finally, to confirm that the posterior hypoblast is an inducer of primitive streak formation, we grafted a posterior hypoblast anteriorly, which resulted in the induction of an ectopic primary axis in 6/6 cases (Supplementary Figure 6A, Figure 4A). This ectopic graft induced a decrease in

**Figure 4:**
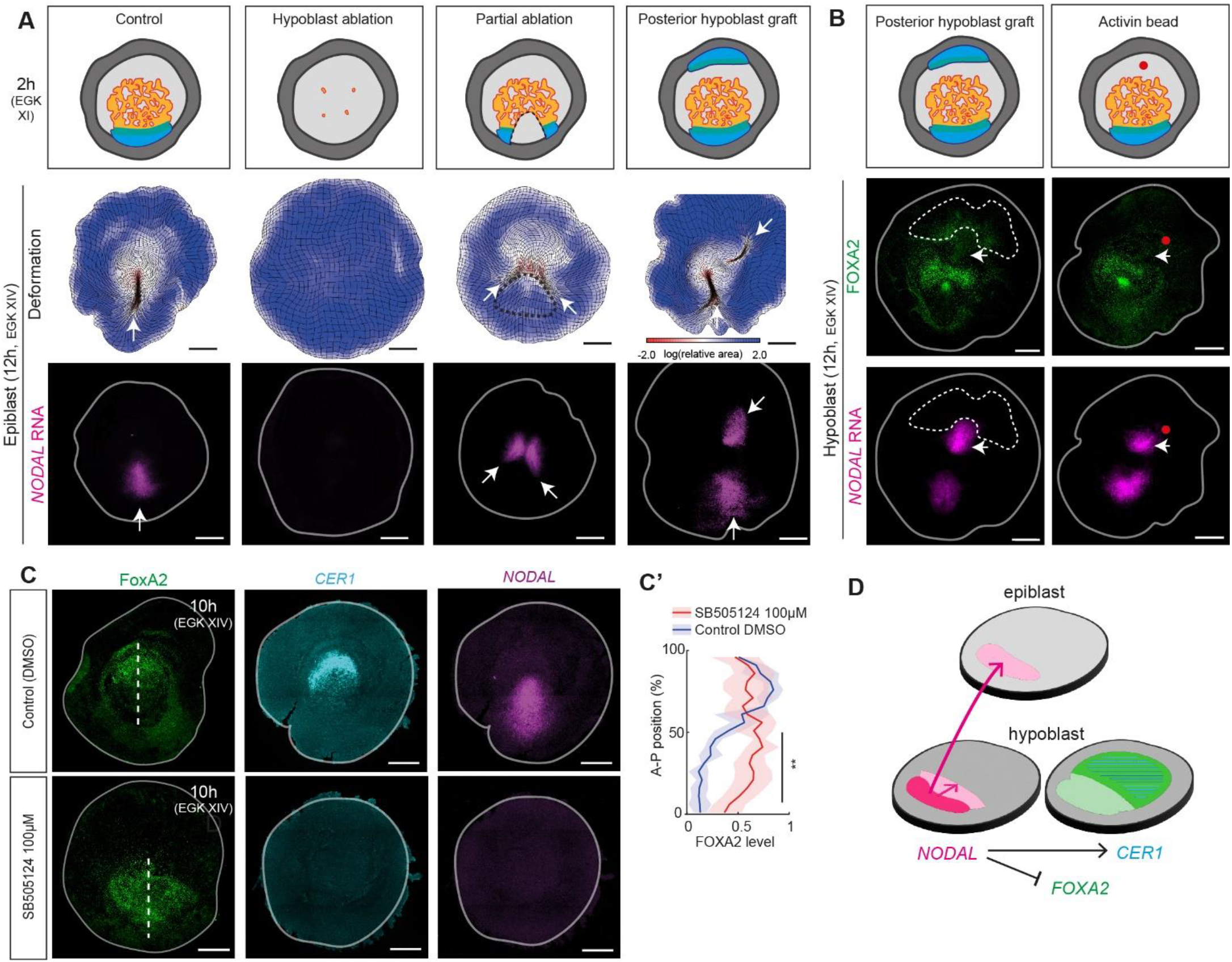
NODAL signaling in the posterior hypoblast induces the primitive streak. **(A)** Top: Diagram illustrating the performed microsurgeries. From left to right: no ablation (control), hypoblast ablation, hypoblast partial ablation, anterior graft of posterior hypoblast. Middle: epiblast deformation maps between 2h-12h for each microsurgery. Bottom: *NODAL* RNA expression in the epiblast at 12h, for each microsurgery. White arrows show primitive streaks (marked by tissue contraction and *NODAL* expression). **(B)** Top: Schematic illustrating an anterior graft of a posterior hypoblast (left) and the deposition of an activin-coated bead anteriorly (right). Middle: *NODAL* RNA expression in the hypoblast at 12h. Bottom: Immunofluorescence for FOXA2 in the hypoblast at 12h. Dotted lines indicate the position of the graft, red dot indicates the position of the grafted bead, white arrows indicate the induction of *NODAL* expression and low FOXA2 levels in the hypoblast. **(C)** Immunofluorescence for FOXA2 and FISH for *CER1* and *NODAL* RNA in hypoblast of control (DMSO) and SB505124-treated embryos at 10h. **(C’)** Averaged FOXA2 levels (+/-std) along the anteroposterior axis (marked by the dotted white lines in c) in the hypoblast of control or SB505124-treated embryos (n=4 control, n=9 SB505124-treated, black line indicates A-P positions for which Welch test p-values < 0.01). **(D)** Graph summarizing the inductive effect of the posterior hypoblast on the surrounding epiblast and hypoblast (light pink: induction of *NODAL* expression, light green: downregulation of FOXA2, dashed blue lines: induction of *CER1* expression). Scale bars: 1 mm.

FOXA2 levels and an ectopic *NODAL* activation in the hypoblast of the host embryo (Figure 4B left panels, n=6/6). Similar results on primitive streak formation and *NODAL*/FOXA2 levels were obtained by depositing a bead coated with Activin A, a known surrogate of NODAL activity^41^, on the hypoblast (Figure 4B right panels, n=7/7). On the contrary, inhibition of NODAL receptor by systemic SB505124 treatment prevented primitive streak formation as well as FOXA2 downregulation and *CER1* expression in the hypoblast (Figure 4C-C’). Taken together, these results demonstrate that NODAL signaling mediates the inducing activity of the posterior hypoblast and that it concomitantly patterns the hypoblast along its antero-posterior axis by shaping *FOXA2* and *CER1* expression domains (Figure 4D).

### The anterior hypoblast is unable to inhibit primitive streak formation

The anterior hypoblast has been proposed to inhibit primitive streak formation through the activity of *CER1*^13^. Although it has been shown that ectopic expression of *CER1* displaces the site of primitive streak formation^13^, whether the anterior hypoblast inhibits primary axis formation has not been directly tested. However, as explained above, we could only detect a clear *CER1* expression after primitive streak formation has initiated, and only in the anterior hypoblast (Figure 3E). To test whether undetectable levels of *CER1* expression or other inhibitors possibly expressed in the hypoblast can inhibit primitive streak formation, we grafted fluorescent anterior hypoblast cells from a 2h-donor onto the posterior pole of a WT 2h-host and monitored effect on primitive streak formation and *NODAL* expression in the host after 10h (Figure 5A). Not only did we observe that grafts do not inhibit *NODAL* expression in the posterior epiblast, but on the contrary, we found that *NODAL* expression and FOXA2 downregulation were induced in the grafted hypoblast (Figure 5A, Supplementary Figure 7A-B). We thus conclude that the anterior hypoblast is unable to inhibit the *NODAL*-mediated induction of primitive streak at 2h. To check whether this inhibition could be effective later on, when the anterior hypoblast starts to express *CER1* (Figure 3E), we performed similar grafts of 7h-anterior hypoblast onto the posterior side of a 7h- or a 2h-host (Figure 5B and Supplementary Figure 7C). These homochronic and heterochronic grafts show that although the 7h-anterior hypoblast does express *CER1*, it is unable to inhibit *NODAL* expression, but instead becomes patterned into a posterior hypoblast, as revealed by *NODAL* and *FOXA2* expression levels. Altogether, these results show that NODAL signaling in the posterior blastoderm overcomes inhibitory signals produced by the anterior hypoblast. To test whether *NODAL* induction in these grafts arises from interaction with the epiblast or hypoblast host, a posterior 7h-epiblast fragment (without hypoblast) expressing *NODAL* was grafted onto the anterior pole of a 2h-host, where the hypoblast expresses *FOXA2* but not *NODAL*. The grafted epiblast induced the formation of an ectopic primitive streak, but importantly induced *NODAL* expression and FOXA2 downregulation in the host hypoblast (Figure 5C, Supplementary Figure 5D), unlike controls, in which an anterior epiblast fragment was grafted anteriorly (Supplementary Figure 7E). Altogether, these results reveal a positive feedback loop from the epiblast to the hypoblast, maintaining NODAL activity in the posterior blastoderm — an effect that, on its own, is sufficient to pattern the hypoblast along its anteroposterior axis (Figure 5D).

**Figure 5:**
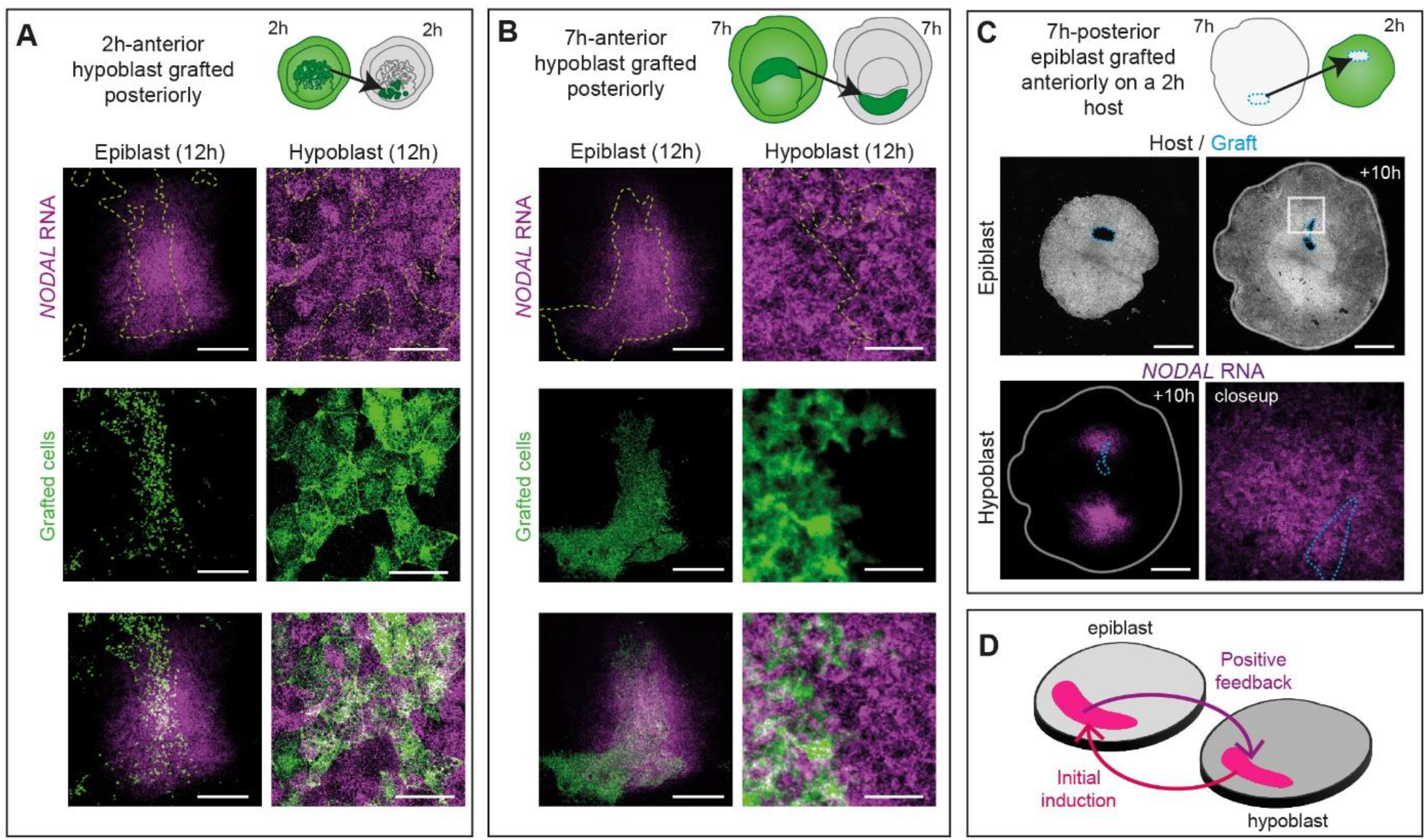
The anterior hypoblast is unable to inhibit primitive streak formation. **(A)** Top: Schematic illustrating anterior graft of transgenic hypoblasts in the posterior of a WT host at 2h. Bottom: Epiblast (left) and closeup hypoblast images (right) at 12h of the obtained chimera. Note the unaffected *NODAL* expression in the epiblast and the induction of *NODAL* expression in the grafted cells. **(B)** Schematic illustrating the grafting of anterior hypoblasts onto the posterior side of a non-fluorescent host at 7h. Epiblast (left) and closeup hypoblast images (right) at 12h of the obtained chimera. Note the absence of effect of the grafts on *NODAL* expression in the epiblast and the induction of *NODAL* expression in the grafted cells. **(C)** Top: Schematic illustrating the grafting of a WT 7h-posterior epiblast onto the anterior side of the epiblast of a memGFP transgenic 2h-host. Middle: Epiblast view of the obtained chimera right after grafting (left) and 10h after graft (right). Dotted light blue contour: grafted tissue. Bottom: Hypoblast view 10h after graft of the obtained chimera stained for *NODAL* RNA and closeup on the anterior side. Note the induction of *NODAL* expression in the hypoblast neighboring the grafted epiblast (n=5/5 chimeras). **(D)** Schematic diagram illustrating the positive feedback loop on *NODAL* expression between the hypoblast and the epiblast after initial induction of *NODAL* expression by the posterior hypoblast. Scale bars: 500 µm (A left, B left), 50 µm (A right, B right), 1 mm (C).

## Discussion

In this study, we show that the original radial symmetry of the blastoderm is broken in the posterior hypoblast, through the localized activation of *NODAL* expression. The propagation of *NODAL* expression induces the formation of the primitive streak in the epiblast and patterns the hypoblast along the anteroposterior axis, repressing *FOXA2* posteriorly and activating *CER1* anteriorly. In the epiblast, *NODAL* induces the large-scale tissue flows leading to primitive streak formation, which concomitantly propagate to the hypoblast through mechanical coupling. Thus, the motion of the hypoblast and its patterning are a consequence and not a cause of primitive streak induction, at odds with the current model in avians and its generalization to amniotes.

In avians, the anterior motion of the hypoblast was proposed to be driven by the intercalation of the endoblast in the posterior, that would displace the hypoblast anteriorly^16,17,19,27,42^. Our data, capturing for the first time the dynamics of the hypoblast at both cell and tissue scales, demonstrate that the hypoblast forms by island fusion and epithelialization without an anterior displacement of the hypoblast. Although we cannot rule-out the contribution of radial intercalation during the epithelization process (between 2-7h, cells rearrange extensively), we do not observe in our dynamic imaging any displacement of the hypoblast relative to the epiblast preceding primitive streak induction. Instead, we show that the hypoblast passively deforms through the transmission of the forces generated in the epiblast that shape the primitive streak^14,29,32^ while the hypoblast itself only shows an intrinsic but modest area expansion. Interestingly, it was previously reported that the extracellular matrix at the interface between the epiblast and hypoblast also exhibits counter-rotating flows^43^, as observed in the epiblast and the hypoblast. It is thus possible that the extracellular matrix may act as a link that transmits active forces from the epiblast to the hypoblast.

Because the hypoblast is entrained and deformed by the epiblast, it remains in constant contact with it during primitive streak induction and formation. Thus, there cannot be a relief of inhibition and indeed an inhibitory function of the hypoblast at all during this process. Consistent with this, we find that expression of *NODAL*, a critical factor in primitive streak formation that was thought to be downstream of *GDF1*^16,36^, initiates in the posterior hypoblast and subsequently propagates and increases in both the epiblast and hypoblast. This propagation of *NODAL* expression is reminiscent of that observed in patterned human embryonic stem cell colonies, in which *NODAL* expression propagates through a short-range relay mechanism^44^. Furthermore, our analysis of the temporal evolution of *NODAL* and *GDF1* expression indeed suggests that *GDF1* is downstream of *NODAL* during primitive streak induction. Such a view is consistent with the recent finding that *PITX2*, a known target of *NODAL* in other developmental contexts^45^, regulate *GDF1* expression during primitive streak induction^46^ and also with the finding that, in other species, GDF1 is a cofactor of NODAL that might be inactive on its own^47,48^. Importantly, surgical ablation of the hypoblast performed in this study prevented primitive streak induction, while anterior grafts of posterior hypoblast induced ectopic primitive streak, confirming the inducing activity of the hypoblast. It is worth noting that similar experiments have been published, but their results were however not mutually consistent. On the one hand, the inhibitory role of the hypoblast was concluded from surgical ablation of the hypoblast, since this procedure occasionally led to the induction of multiple axes^13^. Here we show that supernumerary axes form when isolated pieces of posterior hypoblast are intentionally left behind during surgical ablation. As expected for a local induction process, each of these isolated hypoblast regions induces a primitive streak in the overlying epiblast, resulting in the formation of multiple axes. We note that the complete surgical ablation of the hypoblast is particularly difficult to achieve at these early stages, when the hypoblast is composed of non-epithelialized islands. We therefore controlled that most if not all hypoblast cells were removed following surgical ablation using FOXA2 immunofluorescence and *NODAL* HCR-RNA-FISH (Supplementary Figure 6A). On the other hand, the inductive power of the posterior blastoderm has been identified by various series of grafts, but whether the inductive region was located in the epiblast or in the hypoblast was unclear^11,20,42,49^. Callebaut et al. performed grafts of posterior hypoblast (i.e. Koller’s/Rauber’s sickle) onto explants of central epiblast, which resulted in ectopic primitive streak induction^50^. Here, we obtain similar results and further show that NODAL activity mediates the inducing activity of the posterior hypoblast. Finally, although it is possible that NODAL inhibitors expressed in the anterior hypoblast might act to restrict the activity of *NODAL* and prevent the formation of ectopic primitive streak later on, as observed in mouse^1^, such inhibition of the hypoblast is unlikely to play a role in positioning the endogenous primitive streak, since *CER1* is not expressed early on and the anterior hypoblast is unable to inhibit endogenous *NODAL* activity when grafted posteriorly.

The progressive anterior relocation of a series genes including *NODAL* inhibitors and *FOXA2* in the hypoblast^20,21,22,23^ has been attributed to the displacement of the hypoblast^13,21^. We show that the regionalization of *FOXA2* and *CER1*, and presumably other genes (e.g. *DKK1, CRESCENT*, etc.) is not established by the emergence and invasion of a new endoblast population as previously proposed, but by a genetic regulation induced by *NODAL* activity within the hypoblast. Indeed, we find that *CER1*, which was proposed to initially inhibit NODAL signaling^13^, is expressed at undetectable or very low levels in the hypoblast at earliest stages. Transplantation experiments further show that the anterior hypoblast is unable to inhibit primary axis formation in the posterior, but instead becomes itself patterned into a posterior identity, as revealed by *NODAL* induction and *FOXA2* downregulation. Furthermore, transplanting posterior epiblast fragments from 7h-donners, in which *NODAL* expression has been initiated, in anterior epiblasts shows that the anterior hypoblast of the host downregulates *FOXA2*, an effect which is mimicked by the deposition of Activin A beads. These results place the patterning of the hypoblast concomitant to, if not downstream of, primitive streak formation.

More generally, our results show that primitive streak initiation is positioned by the localized activity of an inducer (NODAL) and not by a relief of inhibition. This result, which redefines how primary axis formation is regulated, has several important implications. First, as early avian development is regulative, and multiple primitive streaks can be induced along the embryo margin upon mechanical perturbations of the epiblast^51,52^, it could be conceived that a primitive streak can spontaneously self-organize at any location along the margin, but that this potential is initially and homogenously inhibited^38,51,53^. However, experiments in which the hypoblast is surgically removed show that this is not the case: in the absence of an inducing trigger (NODAL from the hypoblast) the epiblast cannot self-organize any (single or multiple) primitive streak at its margin. This further suggests that the self-organizing property of the embryo margin^52^ requires at least permissive, if not instructive signals from the hypoblast. Second, our study traces back the first molecular event breaking the original radial symmetry of the blastoderm to the localized expression of *NODAL* in the hypoblast, begging the question of the mechanisms responsible for this restricted expression. Embryonic polarity has been reported to be fixed prior to oviposition, during intrauterine development, under the influence of gravity^54^. How gravity eventually results in *NODAL* expression specifically in the posterior hypoblast remains to be examined. Finally, in mice, several studies have shown that molecular asymmetries in the VE precede its motion and it has been proposed that they predetermine its direction^4,5,6,56,57^. Our results in avian embryos are consistent with the view that changes in gene expression breaks the radial symmetry of the embryo prior to hypoblast/VE directional motion. The recent identification of counter-rotating flows associated with the anterior movement of the VE in mice^55^ is intriguing and — in the light of our findings — should encourage the characterization of the tissues flows occurring in the epiblast and their possible role on the anterior motion of the VE.

## Acknowledgements

We are grateful to Arthur Michaut, Alexander Chamolly and Carolina Parada for their advice on experiments and data analysis. Work in J.G. lab is funded by the European Research Council (ERC) under the European Union’s Horizon 2020 research and innovation programme (grant agreement n° 866186 to J.G), the Agence Nationale de la Recherche (LabEx Revive), the Centre National de la Recherche Scientifique (CNRS) and Institut Pasteur. Y.I was supported by a stipend from the Pasteur - Paris University (PPU) International PhD program and by 4th-year PhD fellowship from the Fondation pour la Recherche Médicale (FRM). For the purpose of open access, the authors have applied a CC-BY public copyright license to any Author Manuscript version arising from this submission.

## Author contributions

The project was conceptualized by A.V. and J.G. A.V. and J.G. designed experiments. A.V. and Y.I. performed experiments. A.V and F.C. designed quantitative image analyses approaches. A.V. and J.G. analyzed the data. A.V. and J.G. wrote the manuscript and generated the figures with inputs from F.C. O.A-P. and C.P. generated the tdTomato:Myosin transgenic quail line and managed transgenic egg supply.

## Materials and Methods

### Animals

All experimental methods and animal husbandry for transgenic quails were performed in accordance with the guidelines of the European Union 2010/63/UE, approved by the Institut Pasteur ethics committee authorization #dha210003, and under the GMO agreement 2432.

### Imaging

#### Hypoblast imaging

Transgenic quail eggs were collected after oviposition and placed in a 38.5°C incubator for 2h to facilitate embryo collection without damaging the hypoblast. Embryos were then collected (stage EGK XI) using a paper filter ring and cultured on a semi-solid nutritive medium made of thin chicken albumen, agarose (0.2%), glucose and NaCl, as described previously^27^. The lid of a 35mm Petri dish was filled with a more solid nutritive medium (with 0.4% agarose), and an embryo was transferred on it, with its epiblast surface and vitelline membrane facing the semi-solid nutritive medium. The lid with the embryo was then inverted onto a glass-bottom dish (Mattek) containing silicone oil (Sigma-Aldrich 378321). This way, the embryo is in contact with the silicone oil on its hypoblast side, enabling confocal live imaging through the coverslip. Embryos were imaged at 38.5 °C using a Zeiss LSM 900 or LSM 980 microscope and 20× or 10× objectives. The time interval between two consecutive frames was ranging from 6-20 min.

#### mEOS2 photoconversion and imaging

Transgenic mEOS2 quail eggs were collected 2h after oviposition and placed on a glass-bottom dish filled with semi-solid nutritive medium. The dish was placed upside down, without lid on a Zeiss LSM 980 microscope with a focus on the hypoblast. 150×150µm square regions were photoconverted at different locations of the hypoblast using a 10× objective. Both epiblast and hypoblast cells were photoconverted in these regions. A photoconversion calibration curve was calculated before each imaging session, and the amount of blue light giving 95% of maximal photoconversion was chosen for subsequent photoconversion. Embryos were imaged (both red and green emission) from their hypoblast and then epiblast sides just after photoconversion and 6h later, using a 10× objective. Fluorescent signal from the epiblast and hypoblast were projected and digitally aligned with each other using the transmitted light image of the embryo, which was collected during the imaging of the embryos.

#### Simultaneous imaging of hypoblast and epiblast using a chimera

Hypoblast cells were collected from a 2h embryo expressing tdTomato:Myosin and grafted onto the hypoblast of a host expressing memGFP. The resulting chimera was placed on a glass-bottom dish (Mattek) and was imaged from its epiblast side, using a Zeiss LSM 980 or LSM 900 microscope and 5× objective, as described previously^26^. The pinhole of the microscope was open wide to collect the red fluorescent signal from grafted cells through the epiblast. The time interval between two consecutive frames was 6-7 min.

### Microsurgery

For all microsurgeries, embryos were collected at 2h after oviposition, and, if needed, incubated for 5h at 38.5°C to reach 7h after oviposition. Before each surgery, the embryo was carefully immersed drop by drop in Hank’s Balanced Salt Solution (HBSS) using a 10 µl pipette. An eyebrow hair mounted on a Pasteur pipette was used to peel off the desired hypoblast tissue (2h hypoblast islands, 2h-posterior hypoblast, 7h-full or anterior hypoblast) or to cut an approximately 500×300µm region of epiblast. Tissues from the donner were transferred in HBSS onto a host using a tapered Pasteur pipette whose tip was cut-off. Each explant was slightly incised on one side before transfer, to ensure proper orientation during the graft.

### Filter intercalation

The hypoblast of a 7h-embryo was first removed and temporarily transferred onto nutritive semi-solid medium. A polycarbonate filter (Millipore RTTP01300) immersed in HBSS, was placed on the epiblast. The hypoblast was then transferred onto the filter using a 10 µl pipette whose tip was cut-off. An eyebrow hair was used to orient and position the hypoblast.

### Pharmacological treatments

#### SB505124 treatment

100µM SB505124 (Sigma-Aldrich S4696, 1% DMSO) and 1% DMSO (control) were added to the culture medium.

#### Activin-coated bead

Heparin-agarose beads (Sigma-Aldrich H6508) were soaked for 2 days at 4°C in a solution containing Fast Green (1:2 dilution, to help visualize the beads) and Activin A (ProteinTech HZ-1138, 50ng/µl). They were then quickly rinsed in HBSS and grafted onto a 2h embryo using a Pasteur pipette.

### Tissue flow analysis

Tissue flows were calculated using Particle Image Velocimetry (PIV) and decomposed into a divergent and a rotational component, as previously described^26^. Spatial alignment of the hypoblast films was performed using the two centers of the counter-rotating flows at 8h as landmarks. Regions masked by debris appearing towards the end of the movies have been filtered out and were not taken into consideration during the analysis. Calculated vector fields were averaged across animals to obtain an average map; regions covered by less than 3 animals were not displayed.

To compare epiblast and hypoblast flows, epiblast flows were analyzed by PIV as previously described^26^, while grafted hypoblast islands were manually tracked. The average hypoblast velocity field was calculated by averaging the velocities calculated over all the animals (providing coverage of the entire hypoblast). The trajectory of each manually tracked hypoblast island was corrected for epiblast movement at each time interval to calculate the movement of the hypoblast relative to the epiblast.

### Immunostaining, RNA-FISH and related image analysis

#### Immunostaining

Embryos were fixed overnight at 4°C in 4% formaldehyde solution. They were then incubated 3×10 min in PBT (1×PBS with 1% Triton 100X and 1% SDS) and overnight in PBT containing rabbit anti-FOXA2 (Proteintech, Cat #22574-1-AP, 1/1000 dilution), or in Can Get Signal Solution B (Toyobo) containing rabbit phospho-Myosin light chain 2 (Cell Signaling, Cat#3674, 1/200 dilution) and mouse anti-ZO1 (Invitrogen, Cat#33-9100, 1/400 dilution). Embryos were rinsed 3×10 min in PBT, incubated for 2h at room temperature with PBT containing secondary antibodies (Invitrogen, donkey anti-rabbit-647 and donkey anti-mouse-555, 1/2000 dilution), for 10 min in PBT containing Hoechst (1/2000 dilution) and for 3×10 min in PBT. They were mounted between two glass coverslips so that both sides (epiblast and hypoblast) can be imaged.

### HCR-RNA-FISH

Embryos were fixed overnight at 4°C in 4% formaldehyde solution and were then labelled using probes specific for quail *NODAL, GDF1* and *CER1* and following the manufacturer’s protocol (Molecular Instruments). The *NODAL* probe was revealed using an amplifier B5-647 or B5-546, the *GDF1* probe using an amplifier B1-546, and the *CER1* probe using B3-647. Immunostainings against FOXA2 were performed after HCR-RNA-FISH labelling.

#### Cryosection

Embryos were embedded in 7.5% gelatin/15% sucrose and sectioned sagittally using a Leica CM3050S cryostat. Sections were degelatinized in PBS at 37°C for 30 min.

#### Imaging and extraction of epiblast and hypoblast signals

HCR-RNA-FISH- and immunolabelled embryos were imaged with a Zeiss LSM 980 or LSM 900 microscope using a 20× or 10× objective (3µm z-resolution) on each side (epiblast and hypoblast sides). Hoechst nuclear signal was used to detect the most superficial plane corresponding to epiblast or hypoblast (depending on embryos’ orientation) and to project fluorescent signals, using custom Matlab code. The projection of the epiblast and hypoblast were then aligned with each other using the contours of the embryo.

#### Spatio-temporal alignment and calculation of archetype maps

Temporal alignment: Embryos used for establishing the evolution of the *NODAL* expression pattern at 4h, 6h and 8h were timed based on the progression of their gastrulation movements, as previously published^49^. Embryos were imaged and their tissue flows calculated on the fly, allowing to fix them at the onset of primitive streak contraction (4h), at 25% medio-lateral primitive streak contraction (6h) or at 50% contraction (8h). Images of the HCR labelling were aligned with the last frame of the live movie using blastoderm contours.

Spatial alignment: the contour of blastoderms was determined at 2h (either directly on the 2h fixed sample, or on the live movie aligned with the staining). A circle was fitted onto blastoderm contours, and its diameter was reduced by 10%. This reduced circle and the position of the posterior pole were used to align all the images (Figure 3B).

Signal normalization: *NODAL* HCR-RNA-FISH signal, which takes the form of a cloud of points each corresponding to the detection of single RNA molecules, was binarized by adjusting a threshold value for each image so as sparse dots were well segmented. *NODAL* expression levels in the epiblast and hypoblast were then averaged for all animals of the same timing.

**Supplementary Figure 1:**
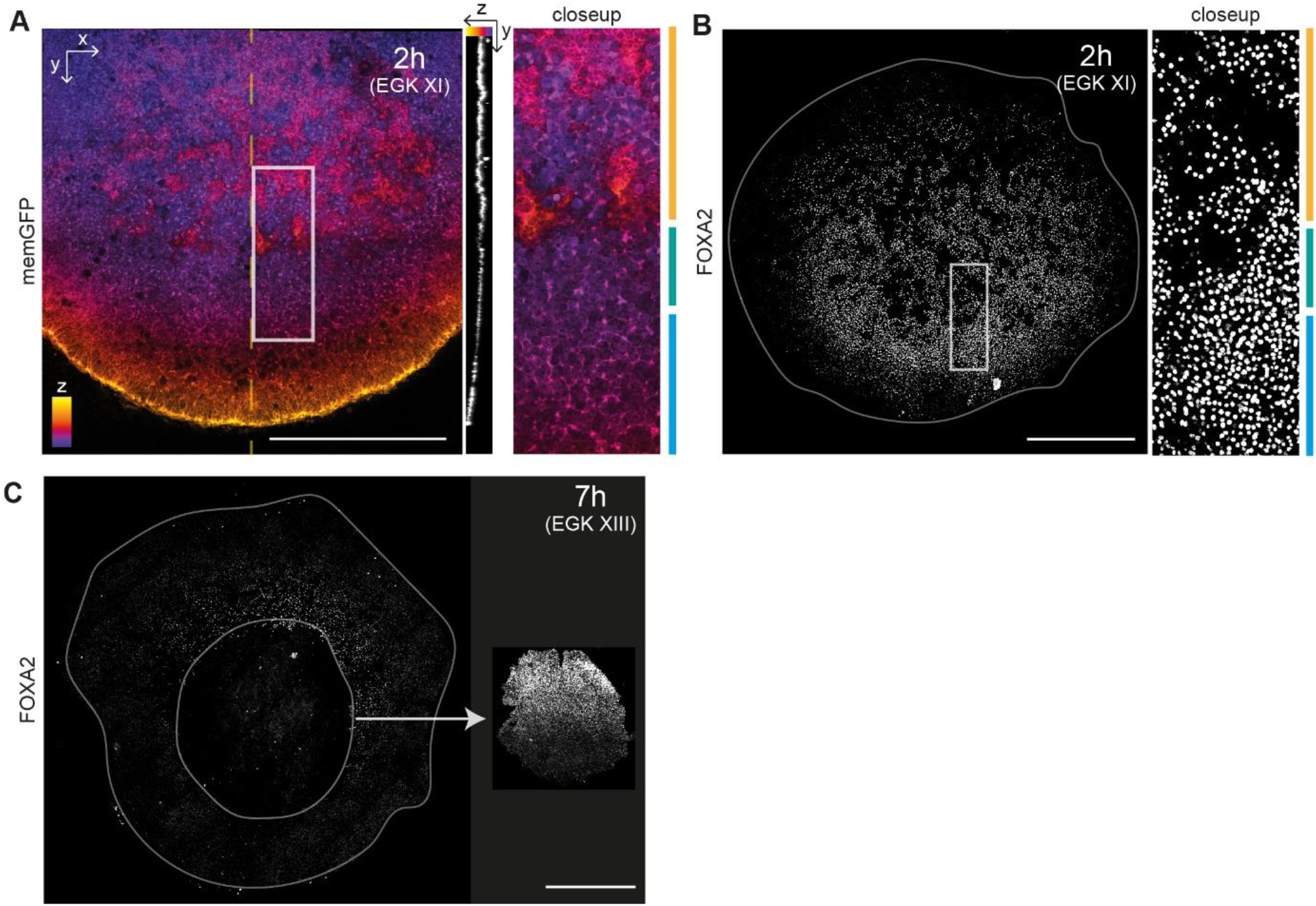
Complementary information on hypoblast imaging and hypoblast populations identification. **(A)** Left: Still image from Movie 1 showing the hypoblast view of an embryo expressing the fluorescent reporter memGFP at 2h. Middle: Optical cross-section along the dotted yellow line. Right: Closeup of the region indicated by the white rectangle. The image is color coded for the Z axis (blue: surface of the deep layers, magenta/orange: superficial layer), highlighting hypoblast islands discontinuity. Orange, green and blue bars, on the right, show the location of hypoblast islands, transition zone and posterior hypoblast respectively. **(B)** Left: FOXA2 immunofluorescence at 2h. Right: Closeup of the region indicated by the white rectangle. Orange, green, and blue bars, on the right, show the location of hypoblast islands, transition zone and posterior hypoblast, respectively. **(C)** FOXA2 immunofluorescence after hypoblast removal at 7h (left: embryo without hypoblast, right: removed hypoblast, n=2 embryos). Scale bars: 1 mm.

**Supplementary Figure 2:**
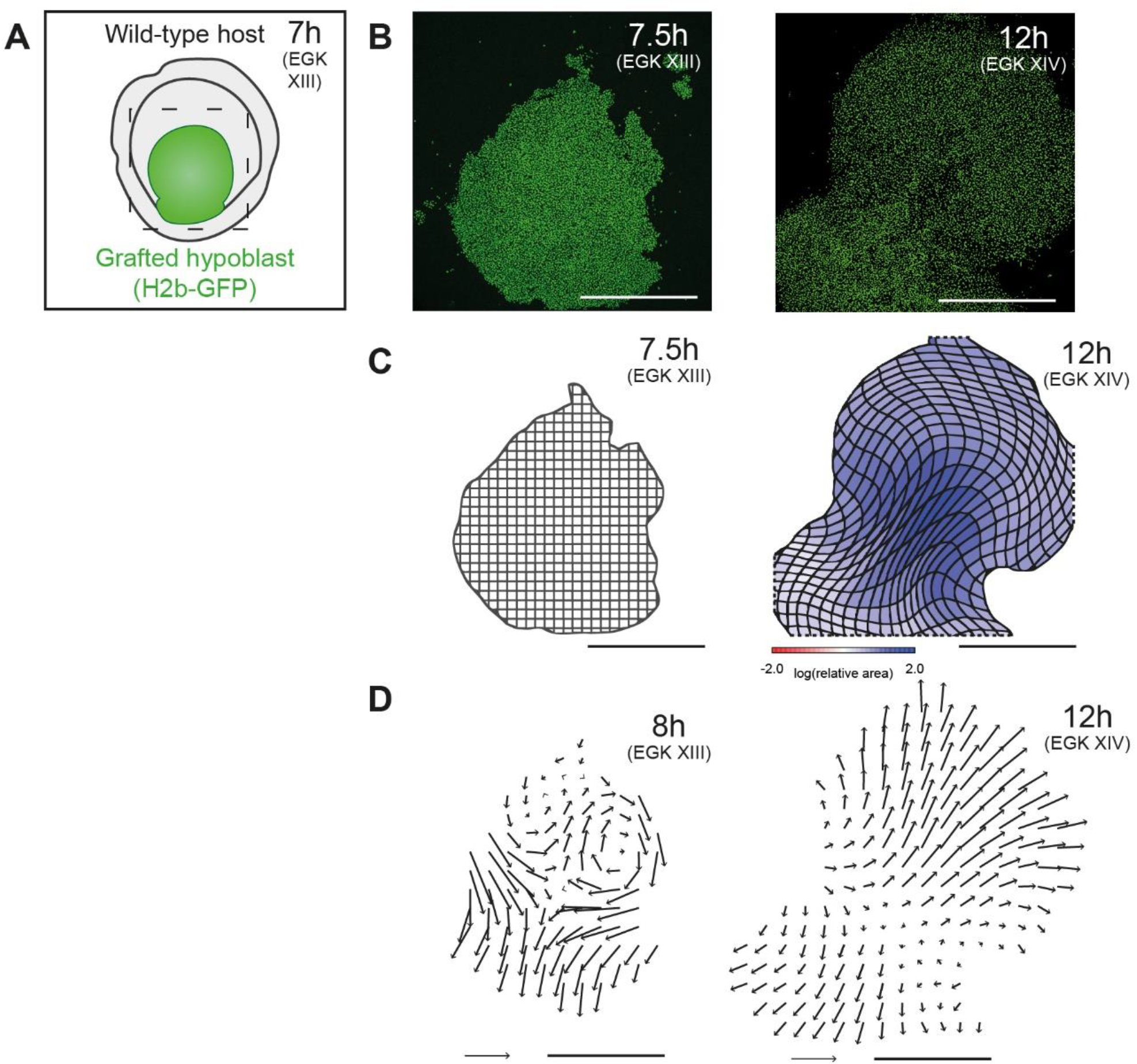
Confirmation of tissue flow monitoring with a chimera. **(A)** Schematic illustrating a chimera obtained by replacing the hypoblast of a 7h wild-type host embryo (gray) with a hypoblast from a 7h embryo expressing H2b-GFP (green). **(B)** Still image of Movie 2 of the grafted hypoblast at 7.5h (30 min after graft) and 12h. **(C)** Cumulated deformation of an initially square grid between 7.5h-12h using PIV. Color code indicates area changes. n=4/4 imaged chimera show similar cumulated deformation. **(D)** Velocity field in the grafted hypoblast at 8h and 12h. n=4/4 imaged chimeras show similar motion fields. Scale bars: 1 mm Scale vector: 100 µm/h.

**Supplementary Figure 3:**
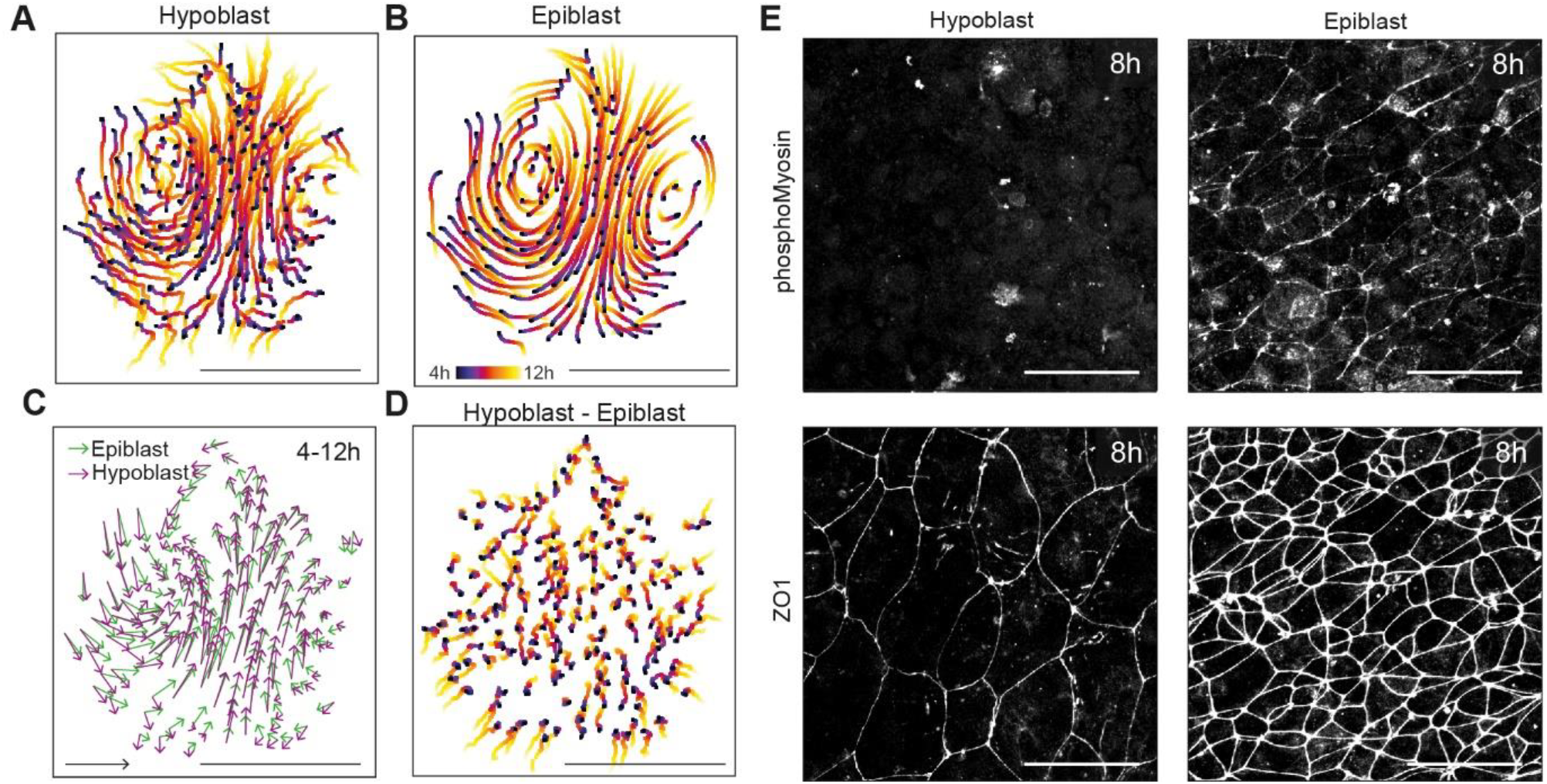
Measure of epiblast and hypoblast flows and phospho-Myosin localization. **(A)** Tracking of tdTomato:Myosin grafted hypoblast cells imaged through the epiblast (158 manual tracking) for a representative chimera. Color code indicates timing (from 4h to 12h). **(B)** Tracks in the corresponding epiblast, obtained by PIV using memGFP signal. Initial points of epiblast tracks match those chosen for hypoblast tracking. **(C)** Averaged velocity vectors between 4h-12h in epiblast and hypoblast for the chimera displayed in (A-B). **(D)** Tracks of the differential motion between the hypoblast and epiblast for the chimera displayed in (A-C). **(E)** Immunofluorescence for phospho-Myosin light chain 2 and ZO1 in the hypoblast and the epiblast at 8h post-laying (absence of cortical phospho-Myosin light chain 2 in hypoblast is observed in n=3/3 labeled embryos). Scale bars: 1 mm (A-D), 50 µm (E). Scale vectors: 100 µm/h

**Supplementary Figure 4:**
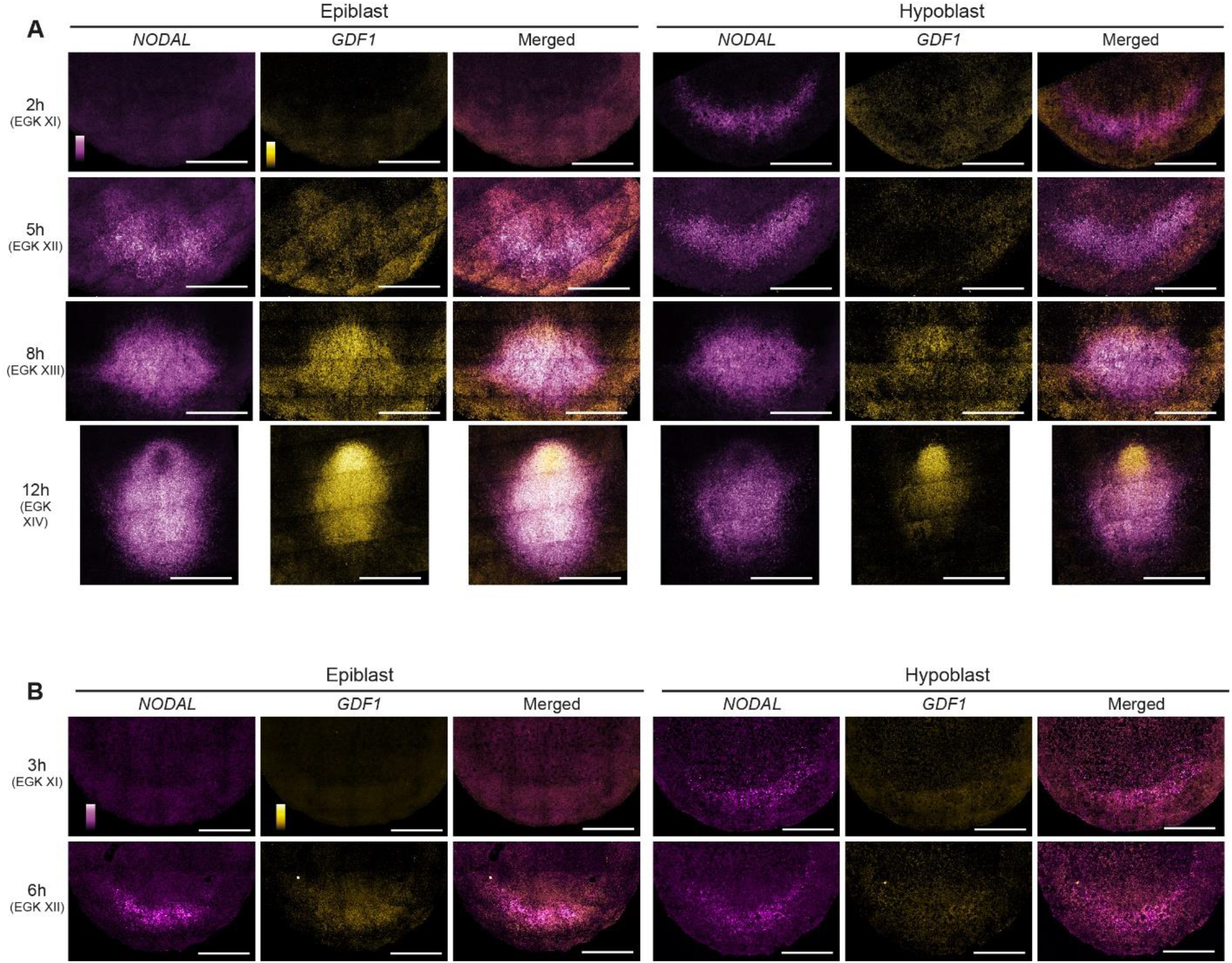
Comparison of *NODAL* and *GDF1* expression patterns using HCR-RNA-FISH. **(A)** *NODAL* (in magenta) and *GDF1* expression (in yellow) at 2h, 5h, 8h and 12h in posterior epiblast and posterior hypoblast of quail embryos. Note the absence of specific signal in epiblast and hypoblast for *GDF1* at 2h (observed in n=9/9 embryos). Note the appearance of *NODAL* and *GDF1* expression in the posterior epiblast at 5h (observed in n=4/4 embryos); and the difference in *GDF1* and *NODAL* expression patterns at 8h and 12h (observed in n=5/5 embryos at 8h and n=9/9 embryos at 12h). **(B)** *NODAL* (in magenta) and *GDF1* expression (in yellow) at 3h and 6h post-laying in posterior epiblast and posterior hypoblast of chicken embryos. Note the absence of specific signal in epiblast and hypoblast for *GDF1* at 3h (observed in n=3/3 embryos). Note the appearance of *NODAL* and *GDF1* expression in the posterior epiblast at 6h (observed in n=3/3 embryos). Scale bars: 1 mm.

**Supplementary Figure 5:**
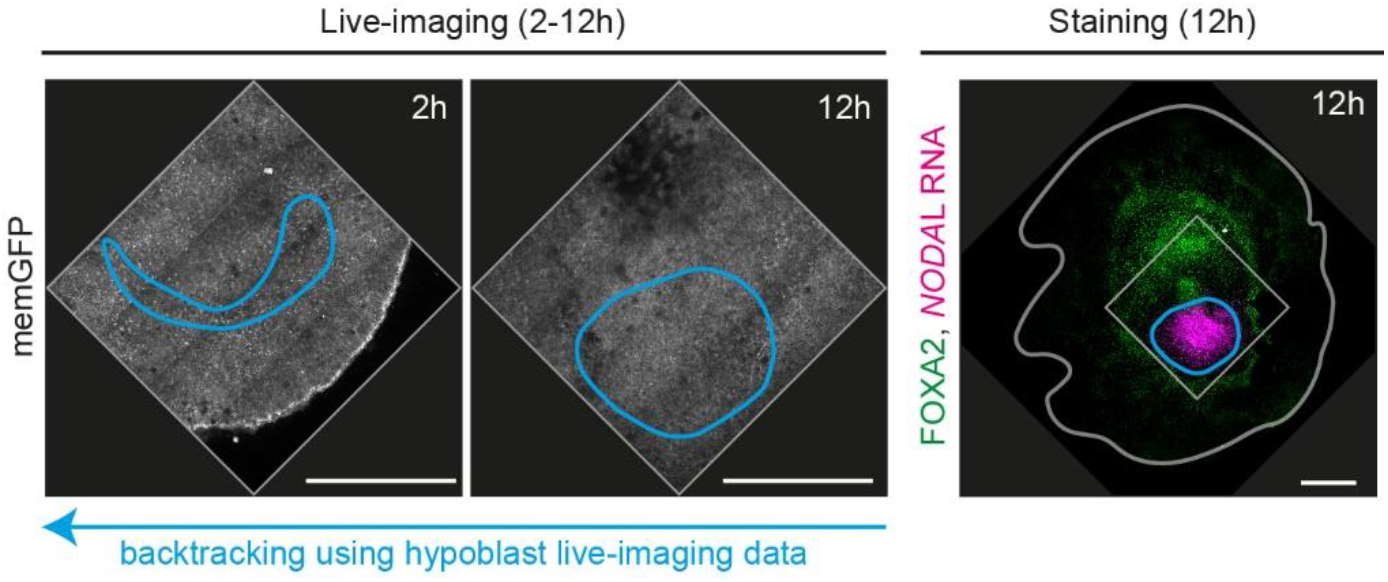
Backtracking the hypoblast population expressing *NODAL* at 12h. Backtracking the *NODAL*-high FOXA2-low region of the hypoblast using hypoblast tissue flows from 12h to 2h. In all of the 4 embryos analyzed, this region is backtracked to the transition zone between hypoblast islands and the posterior hypoblast where FOXA2 is high and *NODAL* expression emerges (see Supplementary Figures 1 and 4). Scale bars: 1 mm.

**Supplementary Figure 6:**
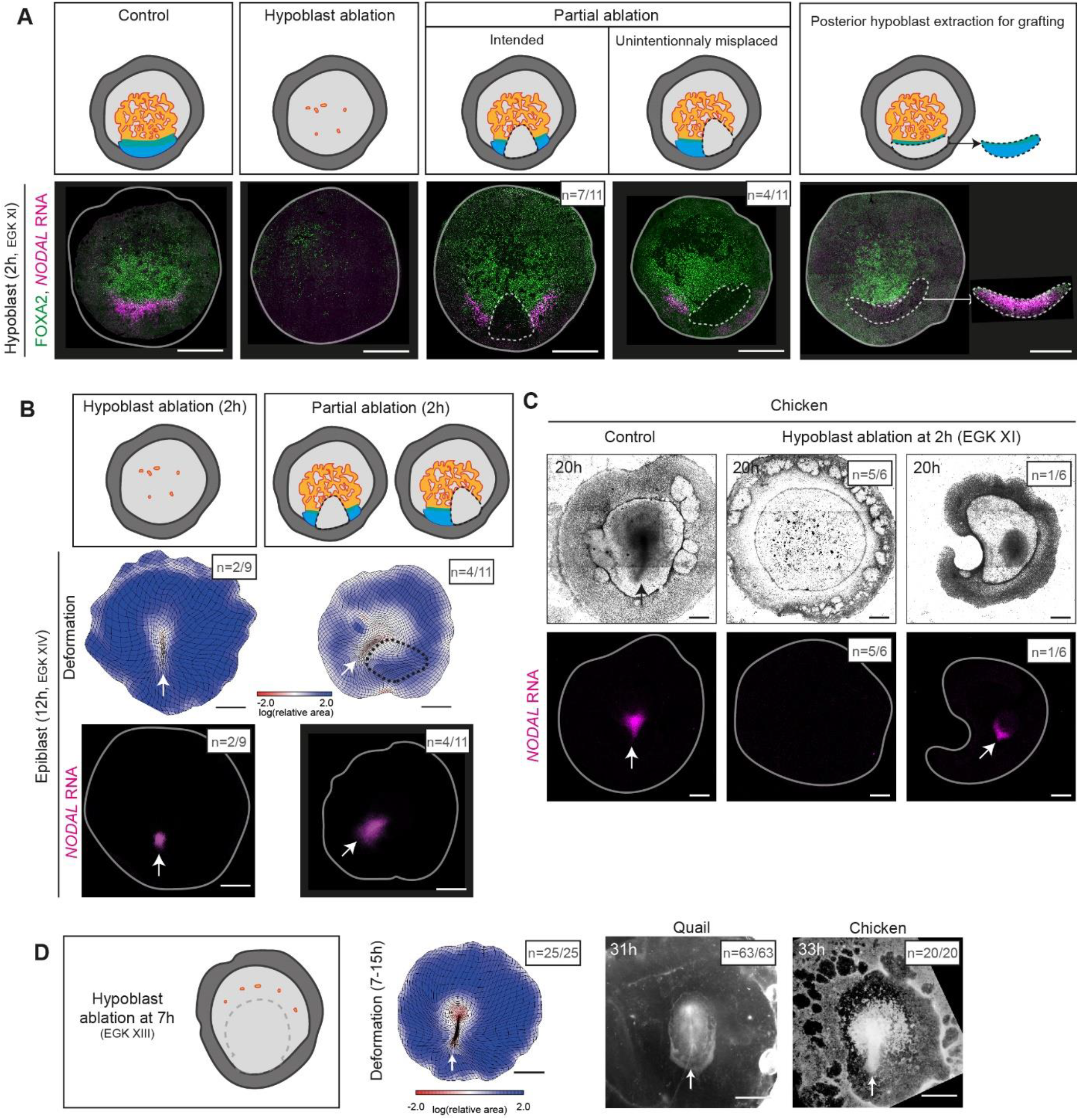
Complementary information on hypoblast microsurgeries related to Figure 4. **(A)** From left to right: schematics illustrating different microsurgeries (from left to right): 2h-hypoblast ablation, partial posterior ablation of 2h hypoblast and 2h-posterior hypoblast extraction, used for grafting on anterior position. Bottom: Verification of *NODAL* expression and FOXA2 levels at 2h for the control and right after surgery for the different microsurgeries. Dashed lines represent ablated region. Note that the precise contours of the posterior hypoblast being unclear under transmitted light, partial and centered posterior ablation can result in one domain of *NODAL* expression (n=4/11) instead of two (n=7/11), as intended by the surgery. **(B)** Infrequent phenotypes observed after 2h-hypoblast ablation (left, n=2/9) or partial posterior 2h-hypoblast ablation (right, n=4/11), complementary to the ones shown in Figure 4. Top: schematics illustrating the different microsurgeries. Middle: epiblast cumulative deformation between 2h-12h (color code represents area changes). Bottom: *NODAL* expression in the epiblast at 12h (white line: embryo contours). White arrows show primitive streaks. **(C)** Top: Transmitted light image in control and 18h after surgery of chicken embryos whose hypoblast was ablated at 2h of incubation. Bottom: *NODAL* expression in the epiblast at 20h (white line: embryo contours). White arrows show primitive streaks. Absence of primitive streak after hypoblast ablation at 2h was observed in 5/6 cases, development of a miniature embryo was observed in 1/6 cases. **(D)** Left: schematic illustrating hypoblast ablation at 7h. Center: Epiblast deformation between 7h-15h in embryos whose hypoblast was ablated at 7h. A single primitive streak was observed in n=25/25 animals. Right: transmitted light image 24h after surgery of quail or chicken embryos whose hypoblast was ablated at 7h, single primitive streak was observed in n=63/63 ablated quail embryos and n=20/20 chicken embryos. Scale bar: 1 mm.

**Supplementary Figure 7:**
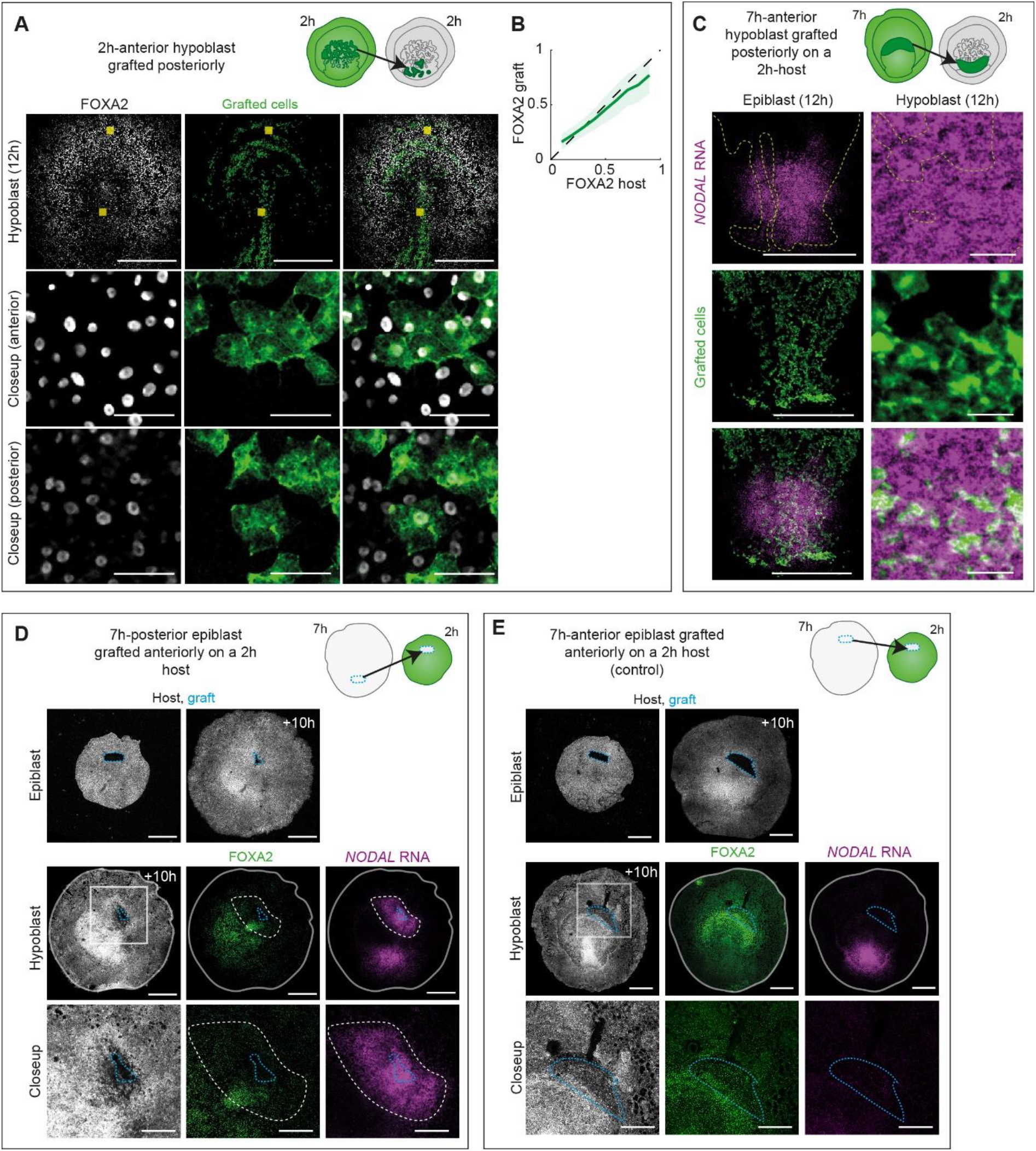
Complementary information on the role of anterior hypoblast and anterior epiblast on primary axis formation, related to Figure 5. **(A)** FOXA2 immunofluorescence at 12h, following an anterior 2h-hypoblast graft on a 2h-host. Top: Schematic illustrating the grafting of fluorescent anterior hypoblast cells onto the posterior side of a non-fluorescent host at 2h. Bottom: Hypoblast of the obtained chimera at 12h stained for FOXA2 (grafted cells are in green). Anterior and posterior closeups (yellow squares in top panels), showing high FOXA2 levels in cells grafted anteriorly, and low FOXA2 levels in cells grafted posteriorly. **(B)** Averaged (+/-std) FOXA2 levels in hypoblast from graft and host (n=5 chimeras) in chimeras obtained by grafting 2h-anterior hypoblast graft onto a 2h-host. **(C)** Top: Schematic illustrating the heterochronic grafting of anterior 7h-hypoblast onto the posterior side of a non-fluorescent 2h-host. Epiblast (bottom left) and closeup hypoblast images (bottom right) at 12h of the obtained chimera. Note the absence of effect of the grafts on *NODAL* expression in the epiblast and the induction of *NODAL* expression in the grafted cells. **(D)** Top: Schematic illustrating the grafting of a non-fluorescent 7h-posterior epiblast fragment onto the anterior epiblast of a fluorescent 2h-host. Middle: Epiblast view of the obtained chimera right after (left) and 10h after the graft (right). Dotted light blue contour: grafted tissue. Bottom: Hypoblast view 10h after the graft stained for FOXA2 and *NODAL* RNA and closeup on the anterior side. Note the downregulation of FOXA2 and induction of *NODAL* expression in hypoblast host cells neighboring the epiblast graft (n=5/5 chimeras). **(E)** Top: Schematic illustrating the grafting of a non-fluorescent 7h-anterior epiblast fragment onto the anterior epiblast of a fluorescent 2h-host. Middle: Epiblast view of the obtained chimera right after (left) and 10h after graft (right). Dotted light blue contour: grafted tissue. Bottom: Hypoblast view 10h after graft of the obtained chimera stained for FOXA2 and *NODAL* RNA and closeup on the anterior side. Note the absence of FOXA2 downregulation and induction of *NODAL* expression in hypoblast host cells neighboring the epiblast graft (n=5/5 chimeras). Scale bars: 1 mm (A top, C top, D top and middle, E top and middle), 50 µm (A closeups, B right, D closeup, E closeup)

**Movie1: Quantitative description of hypoblast dynamics by videomicroscopy**

(1) Live imaging of the hypoblast of an embryo expressing the fluorescent reporter memGFP from to 2h to 12h post-laying. Z is color-coded as in Supplementary Figure 1. (2) PIV-calculated flows. 2 regions tracked using PIV. (3) PIV-calculated flows and deformation corresponding to Figure 1D-D’. (4) Average flows and average decomposition into rotational and divergent components.

**Movie2: Confirmation of hypoblast dynamics with a chimera**

Live imaging of the hypoblast of an H2b-GFP transgenic embryo grafted onto a WT host, and the associated PIV-calculated flows and deformations.

**Movie3: Study of hypoblast and epiblast relative movements using photoconversion**

Photoconversion of square boxes in the blastoderm, imaging of hypoblast and epiblast sides after photoconversion and after 6h. The epiblast of the embryo was live imaged between 2h and 8h to ensure that proper development occurred.

**Movie4: Simultaneous imaging of hypoblast and epiblast movements using a 2-color chimera**

(1) Simultaneous live imaging of the epiblast of a transgenic memGFP host embryo grafted with hypoblast cells expressing tdTomato:Myosin. PIV-calculated deformation of the epiblast and manual tracking of hypoblast cells are displayed. (2) Quantitative comparison of epiblast and hypoblast motions.

**Movie5: Effect of filter intercalation on hypoblast dynamics**

Live imaging of a transgenic H2b-GFP hypoblast grafted on a host embryo with or without filter intercalation, and the associated PIV-calculated deformation grids and tracked circular regions.

**Movie6: Effect of full or partial hypoblast ablation on epiblast dynamics**

Live imaging of the epiblast of a control embryo and embryos in which the hypoblast has been fully or partially removed, and the associated PIV-calculated deformation grids. Transmitted light allows to visualize regions lacking hypoblast as a result of the ablations.

